# An easy-to-assemble, robust, and lightweight drive implant for chronic tetrode recordings in freely moving animals

**DOI:** 10.1101/746651

**Authors:** Jakob Voigts, Jonathan P. Newman, Matthew A. Wilson, Mark T. Harnett

## Abstract

Tetrode arrays are the gold-standard method for neuronal recordings in many studies with behaving animals, especially for deep structures and chronic recordings. Here we outline an improved drive design for use in freely behaving animals. Our design makes use of recently developed technologies to reduce the complexity and build time of the drive while maintaining a low weight. The design also presents an improvement over many existing designs in terms of robustness and ease of use. We describe two variants: a 16 tetrode implant weighing ∼2 g for mice, bats, tree shrews and similar animals, and a 64 tetrode implant weighing ∼16 g for rats, and similar animals.These designs were co-developed and optimized alongside a new class of drive-mounted feature-rich amplifier boards with ultra-thin RF tethers, as described in an upcoming paper (Newman, Zhang et al., in prep). This design significantly improves the data yield of chronic electrophysiology experiments.

## Introduction

There are large application areas in which movable tetrode^22,26,35^ arrays are the method of choice compared to more recent optical methods^6,8^ or higher density silicon probes^15,29,32^. Briefly, the main advantages of tetrode arrays over optical or more recent electrophysiological methods are as follows:

1. They provide the ability to target deep structures that can not be readily accessed optically without removing overlying structures^1,14^.
2. They make it possible to perform recordings from widely distributed structures with custom arrangements of tetrodes.
3. They do not impose limits on recording times. With the drive’s small weight, and using the accompanying headstage and commutator system (Newman, Zhang et al., in prep), arbitrarily long, uninterrupted recordings can be made. This is in contrast with most optical methods that typically suffer from indicator bleaching or overexpression.
4. Drives are adaptable mechanical devices that are compatible with complex free behavior, especially when used with wireless data loggers^28,30^, and/or miniaturized headstages (Newman, Zhang et al., in prep), and can be combined with other devices.
5. Adjustment of depth of individual electrodes is possible at any time. This has a number of advantages, including making it possible to target deep structures that are otherwise hard to hit precisely, via repeated adjustments and identification of electrophysiological markers of the target. Similarly, adjustable tetrodes help in targeting superficial structures that are otherwise hard to pinpoint due to brain deformation during surgery and probe insertion. Depth adjustment is not restricted to tetrode drives and can be combined with laminar probes, either in single drive^5,33^, or multi-drive^23^ configurations.
6. Tetrodes provide good stability of recording conditions over time. Even though the stability of individual neurons cannot be guaranteed over days^11,21^, through repeated small adjustments, tetrode arrays can be stable over around a year^34^. Given appropriate analysis methods, individual neurons can be recorded from for weeks^10,13^. This makes it possible to record during experiments spanning multiple weeks or months, or to record from tetrodes implanted before a lengthy training period. While optical methods can in some cases rival this long term stability, silicon probe implants typically cannot^16,19^, though good long term results can be achieved^27^.
7. The cost of tetrode drive implants is low compared to silicon probes.

The main downsides of tetrode arrays are:

1. The number of simultaneously recorded channels is typically limited to 32 or 64 in mice, and 64 or 128 in rats, due to the physical size of the implants and due to the effort required to build and load drives with high channel counts. This leads to unit yields that are, in most brain areas, smaller than what can be achieved with miniaturized head-mounted one-photon microscopes^1^.
2. The time and expertise needed to build drive implants is significant. Common ways around this are to use static electrode arrays or static silicone probe implants^5,15^, simplified designs with only one drive mechanism^7,20,33^, or to load subsets of drives in existing multi-drive implant designs.

Recent designs (Halodrive, Neuralynx, Bozeman, MT) for use with 4 tetrodes for mice and 10 or 18 for rats lower the number of steps required to build drives by making use of robust 3D-printed drive bodies that unite multiple functions in one part which reduced the complexity of the assembly process and improves mechanical robustness.

Drive designs are chiefly characterized by the method used to enable stable and precise linear motion:

1. Classic designs^18^ use a screw that is moved into a threaded hole in the drive body. A shuttle piece is fixed to the screw but prevented from rotating by means of one or more guide pins, resulting in linear motion.
2. The spring-based Flexdrive design^34^ replaces the guide pins and shuttles with a one-piece leaf spring that is bent down by the screw resulting in linear motion. This reduces the weight of the drive, but introduces some nonlinearity to the screw travel, requires difficult-to-manufacture springs, and makes drive assembly difficult.
3. Guide channel designs (Neuralynx Halodrive, OvalDrive^4^, and Yamamoto et al. 2008^36^) replace the guide pins with 3D printed channels that prevent rotation of the shuttles. This approach requires higher precision 3D printing, but significantly reduces the overall drive complexity. In some designs, the screw moves down, as in a traditional design, while in others, the screw rotates, pushing a threaded shuttle piece down (Halodrive).

The design described in this manuscript combines a guide channel mechanism with a custom screw that eliminates the need for a separate means of screw retention. The resulting drive is fast to assemble, robust, and maintains a ∼2 g weight for 16 movable drives, with a 4.5 mm travel range for mice (Fig. 1A), and a ∼16g weight for 64 drives and ∼10mm travel range for rats (Fig. 1C).

**Figure 1:**
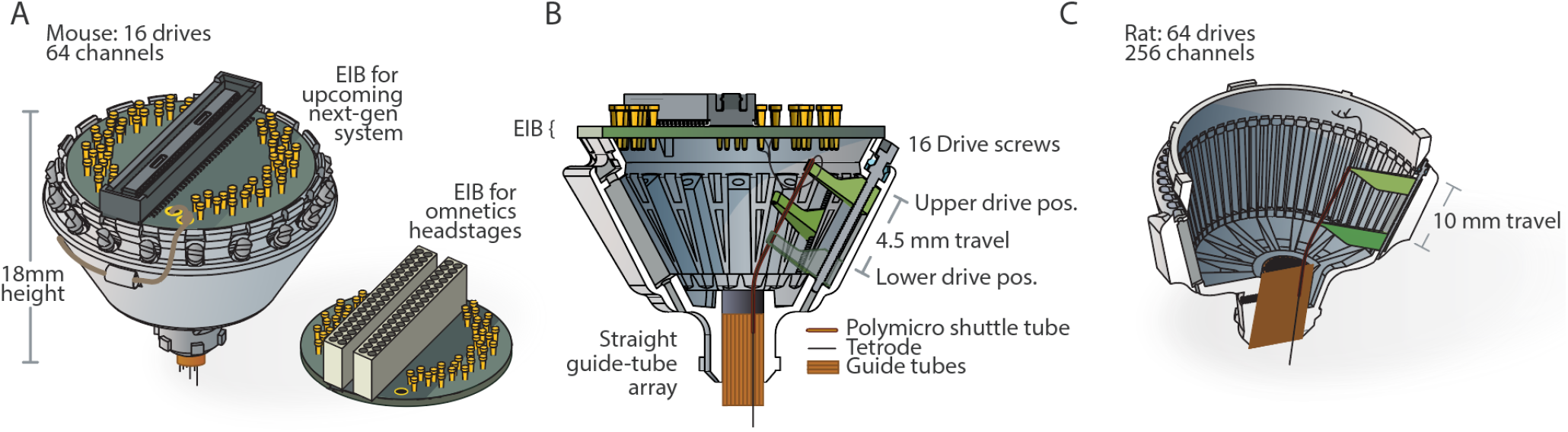
Overview of the drive design. **A**, Overview of the mouse drive implant for 16 individually movable tetrodes and 64 channels, shown with an eletrode interface board (EIB) and the miniature headstage (Newman, Zhang et al., in prep). **B**, Overview of the internal drive mechanism - linearly moving *shuttles* (green) are moved up and down in *guide channels* by captive screws. A straight *guide tube array* (orange) holds fused silica (Polymicro) *shuttle tubes*. **C**, Drive variant with 64 individually movable tetrodes for up to 256 channels, for use in rats, shrews, and similarly sized animals.

A similar, lower density design is currently commercially available from Neuralynx (HaloDrive). This design performs well when 4 tetrodes are sufficient for for mice (or equivalently 10 or 18 tetrodes in rats), and can be used instead of the presented high density design in cases where the high number of concurrently recorded tetrodes is not required.

All design files, parts lists, and drawings are available on the git repository at https://github.com/open-ephys/shuttle-drive. Detailed assembly instructions (and this manuscript) are also available on the repository. This documentation describes Open Hardware and is licensed under the CERN OHL v. 1.2. See the repository at https://github.com/open-ephys/shuttle-drive for a copy of the license.

## Results / Design Overview

Our design can be broken down into two functional components: the *shuttle tubes* and *guide tube* array, and the drive mechanism. See Fig. 2C,D for an overvierw of these components. Each of these two components takes advantage of a technical advancement over the previous flexDrive design: The *shuttle tube* mechanism is enabled by the recent availability of stiff polyimide coated fused silica tubes (Molex Polymicro tubes sn: TSP100170) that provide much higher stiffness than pure polyimide tubes. The drive mechanism is enabled by the availability of high resolution 3D printing that makes it possible to create a sliding friction fit *shuttle* retainer that is sufficiently precise for positioning tetrodes.

**Figure 2:**
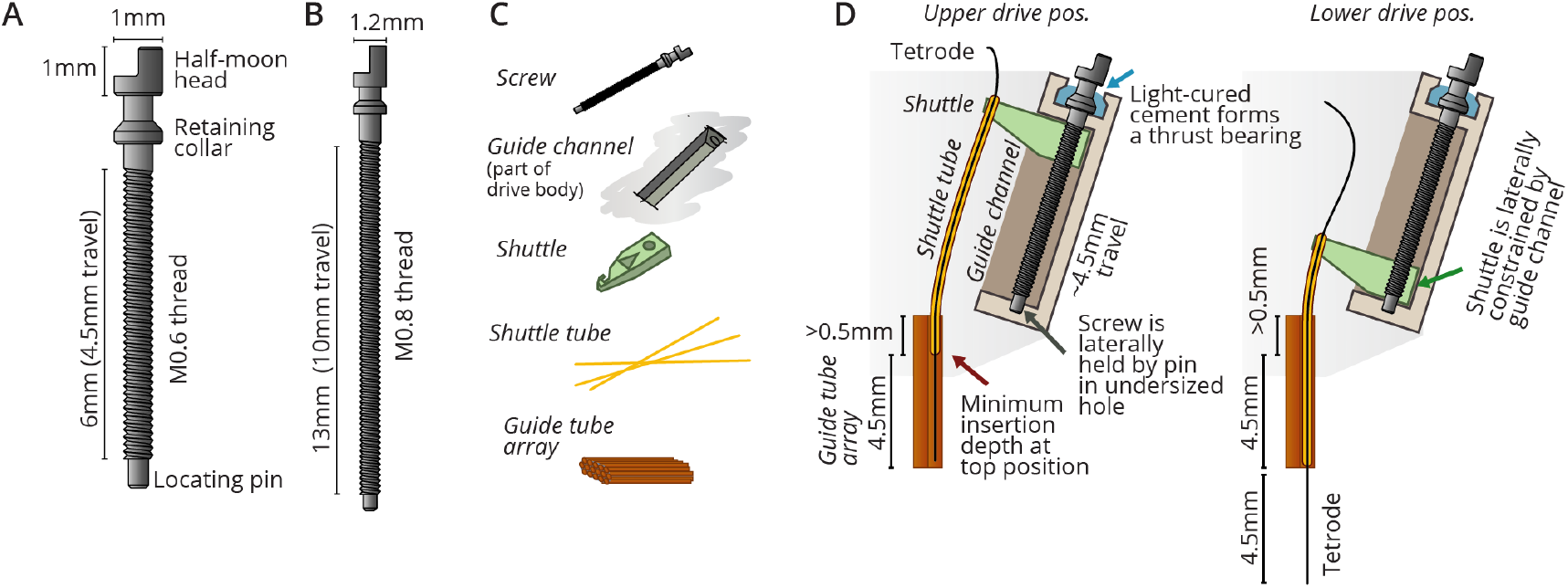
Custom screw design and drive mechanism. **A**, Custom screw for the mouse drive. The central novel features are the retaining collar under the screw head, which acts as a thrust bearing, stopping the screw from moving up, but allowing rotation, and the locating pin at the bottom, which allows the screw to rotate but not move laterally. **B**, Screw variant for use in the 64 drive design for rats, shrews, and similarly sized animals. ∼10mm of travel can be achieved. **C**, Key components that make up one of the linear adjustment (‘drive’) mechanisms. **D**, Overview of the internal drive mechanism - linearly moving *shuttles* (green) are actuated by captive screws. The screws move inside *guide channels* and are held at the bottom via their locating pins, and are held vertically at the top by gluing their retaining collars into concave pockets in the drive body. A straight *guide tube array* (orange) holds fused silica (Polymicro) *shuttle tubes*. At the topmost drive position, the *shuttle tubes* stay inserted in the *guide tube array* - this amount of insertion, plus the desired travel range, dictates the height of the *guide tube array*. The present design achieves ∼4.5mm of travel.

### Drive Mechanism

The drive mechanism is formed by a moving *shuttle* that is pushed up and down by a rotating but otherwise retained screw^36^ (Fig. 2D). The *shuttle* is constrained from rotating along with the screw by the *guide channel* that is part of the drive body.

The screw needs to be prevented from being pushed up and down by the friction of the *shuttle* and/or the force exerted by the adjustment screwdriver. In some previous designs this was accomplished by means of a nut glued to the screw (Neuralynx Halodrive, Yamamoto et al. 2008^36^). Here, a custom screw is used that includes a retaining collar under the screw head that is glued into a recess, providing a rudimentary thrust bearing with a high degree of stability (Fig. 2B). In addition, the bottom of the screw is retained laterally in a press-fit hole in the drive body.

**Table 1:** Design parameter overview for the two default configurations outlined here. Weight will increase by approximately 1.2g (mouse) or 4.7g (rat) when the next-gen headstage (Newman, Zhang et al., in prep) is attached.

|  | Mouse drive | Rat drive |
| --- | --- | --- |
| N drives mechanisms / tetrodes | 16 | 64 |
| N recording channels | 64 | 256 |
| Weight | ~2g, ~(3.5g as implanted,<br>with headstage from<br>(Newman, Zhang et al., in prep),<br>cement, ground wire, etc. | ~16g |
| Max. travel | 4.5mm | 10mm |
| Diameter / Height | 19mm / 14mm | 48mm / 35mm |

The sizing of the drive components are chosen to be as small and closely spaced as possible. M0.6 screws are the smallest screw that is sufficiently robust in the required lengths. In both the mouse and the rat design, the position of the drive mechanisms are governed by spacing constraints that derive from the screw dimensions: the radial distance of the drives from the drive center is determined by the N of drive mechanisms (16 and 64), the width of the *shuttle* (determined by the drive screw size), and the need to maintain some wall thickness between *guide channels*. Additionally, the drive mechanisms are angled outwards by 32° in order to provide an opening for loading *shuttle tubes* into the *guide tube array* when the drives are in a raised position, and in order to provide enough area at the top of the drive to attach the electrode interface board (EIB) and amplifier board. At the bottom position, the *shuttles* are almost touching each other.

### Guide & Shuttle Tube Subassembly

Typical drive (or microdrive) implant designs^2–4,7,20,22,23,25,26,35^ are made up of stereotyped static and moving components to implement tetrode positioning: a static array of tubes, the *guide tube array*, defines the lateral spacing and location of tetrodes (Fig. 1B). Within the static *guide tubes*, moving tubes, called *shuttle tubes* are moved by the drive mechanism. This is done either with a part called a *drive shuttle* (Neuralynx, and^4^), or a spring^34^ that connects the *shuttle* tube to the screw. In almost all designs with > 6 drive mechanisms, the screws and drive mechanisms are arranged at an angle, and the guide tubes fan out from their compact arrangement at the bottom, in order to meet the *shuttle* tubes and drive mechanisms. In classic designs, shuttle tubes are typically made from polyimide, and guide tubes from either polyimide or stainless steel.

The drive design (Fig. 1) uses polyimide coated Silica *shuttle tubes*, which are much stiffer than polyimide. The coating protects the glass and makes the tubes highly resistant to scratching which otherwise makes glass tubes highly fragile. The high stiffness of the tubes allows the shuttles to maintain a long unsupported distance between the *shuttle tubes* and the *guide tubes*, and even bend at limited angles without risking any kinking. This property removes the need for bent individual guide tubes that are otherwise required to ensure a smooth transition of the *shuttle tubes* between the angled drive mechanism axis and the straight guide tube bottom. This approach has been used successfully in the Halodrive design (Neuralynx, Bozeman, MT).

The guide tube assembly can therefore be constructed from a straight extrusion profile, either made from bundles of polyimide tubes, milled from plastic, or possibly even EDM machined. We also tested a variant where we 3D printed the guide tube array in a very high resolution process (stereolithography with around 0.05mm x/y feature size). This 3D printed guide tube array does not provide the same degree of smooth motion as bundled polyimide tubes, and the long-term biocompatibility of the printing material would need to be thoroughly tested before this method is ready for widespread use.

Most drive mechanisms suffer from some amount of hysteresis when reversing the direction of travel. This hysteresis is defined as the amount of applied screw rotation required to take up slack in the mechanism before the tetrode itself starts reversing its motion. In the present design, there are a few different sources of hysteresis: (i) the screw itself can travel up and down relative to the drive body. The method of gluing the screw retaining collar into a pocket on the drive body minimizes this travel, and if glued properly, this hysteresis should be negligible. (ii) If the sizing of the *shuttle* relative to the *guide channel* is not correct, the *shuttle* can change its angle, or move side-to side or even up/down relative to the screw, which can cause some hysteresis. In properly sized *shuttles*, this effect should be negligible. (iii) The slightly bent *shuttle tubes* act as springs, and together with the friction between the *shuttle tubes* and the *guide tube* array, this is the main cause of hysteresis in the drive. This hysteresis is worst for *shuttle tubes* near the center of the *guide tube array*, where the bending of the *shuttle tubes* is maximal, and less near the periphery. Applying lubrication to the *guide tube* array, and making sure that its polyimide tubes are not deformed, can reduce the friction and therefore the hysteresis. See Fig. 3 for measurements of the drive range and hysteresis of the presented drive design. In practice around 75 to 150 μm of hysteresis can be expected for the mouse drive.

In addition to this mechanical hysteresis observed on the drive mechanism in air, another level of biological hysteresis and changes in recorded units are expected from the reaction of the tissue to tetrodes.

**Figure 3:**
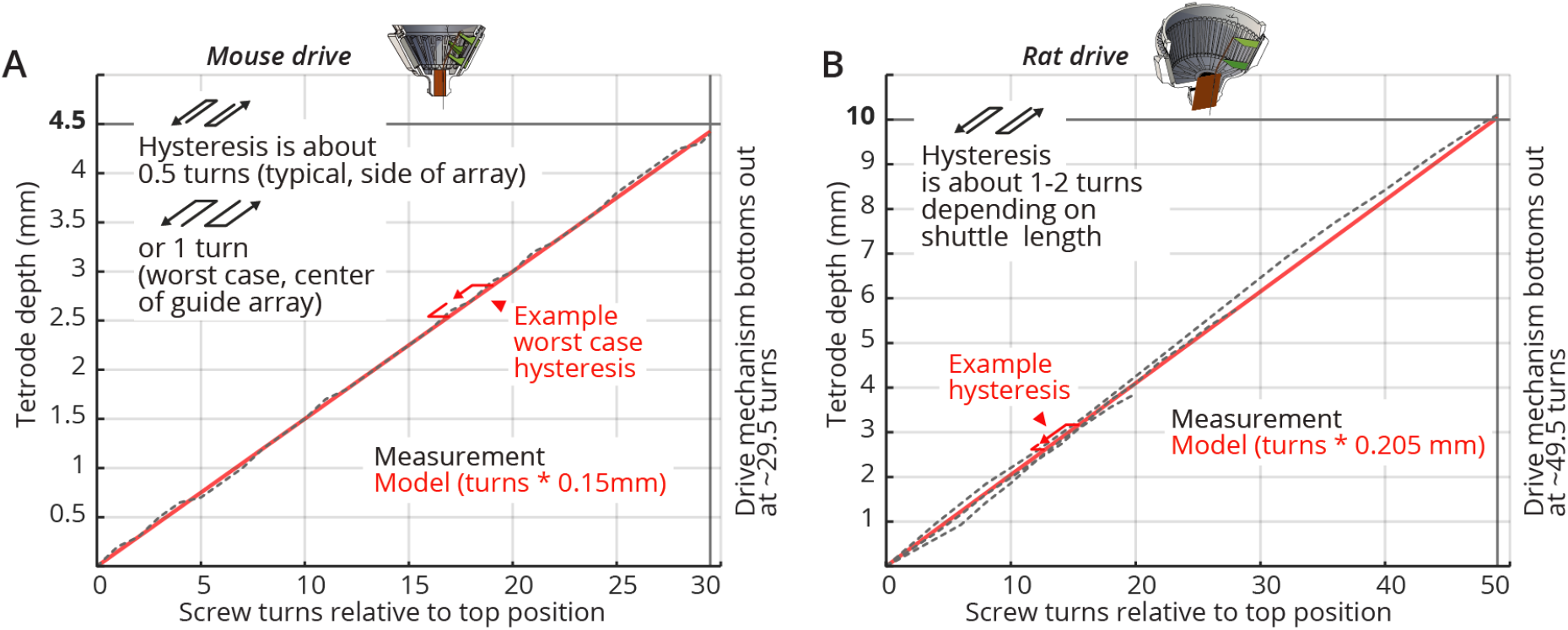
Tetrode depth measurements. Measurements of tetrode depths over the entire adjustment ranges for the mouse (**A**) and rat (**B**) variants. This travel was measured only in the downward direction; some hysteresis will occur when reversing the drives. Hysteresis is measured as the amount of screw rotation after a direction reversal at which point the tetrode started moving again. Hysteresis is caused by different factors; see main text for a short discussion. See inserts for typical measured hysteresis. The mouse drive behaves almost completely linearly with a 1:1 correspondence of screw pitch (0.15mm/turn) to travel range. The rat drive shows some scaling of the travel with a 0.205mm/turn vs. the 0.2mm pitch screw. This factor was measured for a tetrode at the periphery of a circular *guide tube* array, and other arrangements may result in slightly different factors. We recommend individual calibration if this level of dead-reckoning precision is desired. Reaching the design travel ranges of 4.5mm and 10mm requires starting at the absolute top position and moving until the *shuttle* touches the bottom position. Extra care is needed at these positions in order to not drive the *shuttle* into the end stops and strip the threads.

See Dhawale 2017^9^ for a discussion of the expected effects.

### Electrode Interface Boards (EIBs)

The EIB on this design is a variant of the previous EIBs, with the vias for mounting the tetrode wires moved inside from the periphery of the PCB in order to avoid interference between the gold pins and the rim of the drive body. The EIB is attached to the drive body along its entire circumference after tetrodes have been loaded. This shields the entire drive mechanism from outside influences electrically and mechanically, making adjusting the drive safer, and reducing the risk of bedding material and/or dust interfering with the tetrodes.

The drive is compatible with EIBs with classic pinouts that fit common headstages with male omnetics connectors. Other variants can easily be made by modifying the design files supplied in the git repository.

For the mouse variant of the drive, the EIB mounts to the topmost edge of the drive, which has a 17.2mm outer and ∼15.9mm inner diameter. EIB diameters of around 17mm will work best, and significantly larger EIBs run the risk of occluding access to the screws. The number of channels than can be connected on an EIB of this size is mainly limited by the connector footprint, and to a somewhat lesser degree by the available space for gold pins. The holes for the gold pins need to be set back from the edge of the EIB by ∼1.5mm to avoid interference between the gold pins and the angled drive body. Currently, we have designed EIBs for 64 channels, corresponding to 16 tetrodes, in variants for existing classic headstages and for the next-generation miniaturized headstages.

For the rat variant, designs for 128 and 256 channels are available. The size constraint for the number of channels on the EIB applies less in this design. 256 channels for 64 tetrodes are a reasonable compromise in terms of practical experimental considerations and from a point of view of amplifier/headstage design.

### Headstages

The drives can also be used with traditional headstages (Open Ephys, Intan, Neuralynx, Plexon etc.) by using the omnetics EIBs, or with a variety of other systems and headstages by making custom EIBs with an appropriate connector and layout.

## Assembly

The assembly process for the drives is fairly simple compared to previous designs, but differs in some key steps. We recommend that even experienced drive builders thoroughly read the assembly steps when initially learning to build this design. The usual tools - forceps (almost all work on the drive can be done with #3 Dumont forceps or similar), glues (mostly 5min epoxy, and a light-hardened dental cement, etc.), and a stereo microscope are needed. Also, for the final tetrode loading steps, custom jigs are needed in order to hold EIB and drive body at a stable distance to each other while tetrodes are being loaded. See the repository (https://github.com/open-ephys/shuttle-drive) for a comprehensive list of tools and bill of materials (BOM), as well as the section on part availability and manufacturing. Different building methods, variants and instructional videos are linked from the “readme” file on the git repository.

*Additional details:*

Some parts of the build process description are marked as *“Additional details”* and can be skipped for a quicker overview of the method.

### Prepare Electrode Interface Board (EIB)

Solder connector and ground wire as needed. For omnetics-based headstages, regular reflow-soldering or hand-soldering are possible, but for EIBs with more than one connector, extra care must be taken to ensure that the connectors are well aligned in order to allow insertion of the headstage. Relatively small mismatch between connectors can make insertion impossible.

### Prepare guide tube array

The guide tube array can be made similarly as in traditional drive designs^3, 7, 18, 22, 23, 25, 34, 35^ by gluing together layers or bundles of polyimide tubes. The inner diameter (ID) of the tubes needs to fit the outer diameter (OD) of the polymicro tubes (164±06μm), so ∼30ga polyimide tubes will work well (see Parts list and documentation in the repository (https://github.com/open-ephys/shuttle-drive) for manufacturer part numbers and supplier information). The height of the guide tube array needs to be at least the desired travel range (∼4.5mm for mice and ∼10mm for rat drives) plus some allowance for a minimum insertion depth of the polymicro tubes in the guide tube array at the top drive position of around 500 μm(depends on the guide tube diameter and smoothness, etc.). A longer guide tube array does not cause any issues and just makes this minimal required insertion range larger.

Glue longer lengths of tubes into bundles with either superglue or epoxy (Fig. 5A). This can be done in layers, or in bundles.

**Figure 4:**
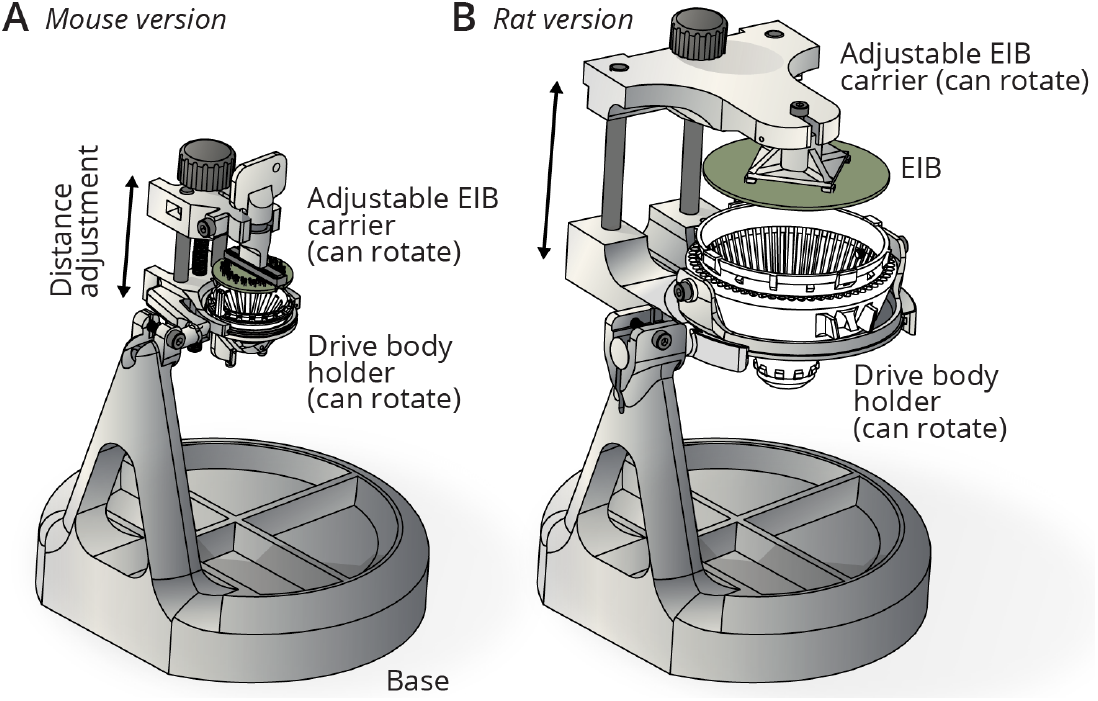
Assembly jigs. **A**, Assembly jig for holding the drive body and EIB in fixed relative positions. The distance between EIB and drive body can be adjusted with a linear slide, driven by a screw. Both the EIB and the drive body can be independently rotated around their principal (vertical) axes by loosening lock screws and manual rotation, in order to facilitate access to all sides of the drive for drive mechanism assembly and tetrode loading. The whole linear slide mechanism which holds the drive and EIB can rotate back and forth (manual, with a friction adjustment) to provide access to the sides of the drive and EIB and to the bottom of the guide tube array (for instance for cutting tetrodes). The jigs are made from easily available stock components and 3D printed custom parts. **B**, Larger jig variant for use with the rat drive.

**Figure 5:**
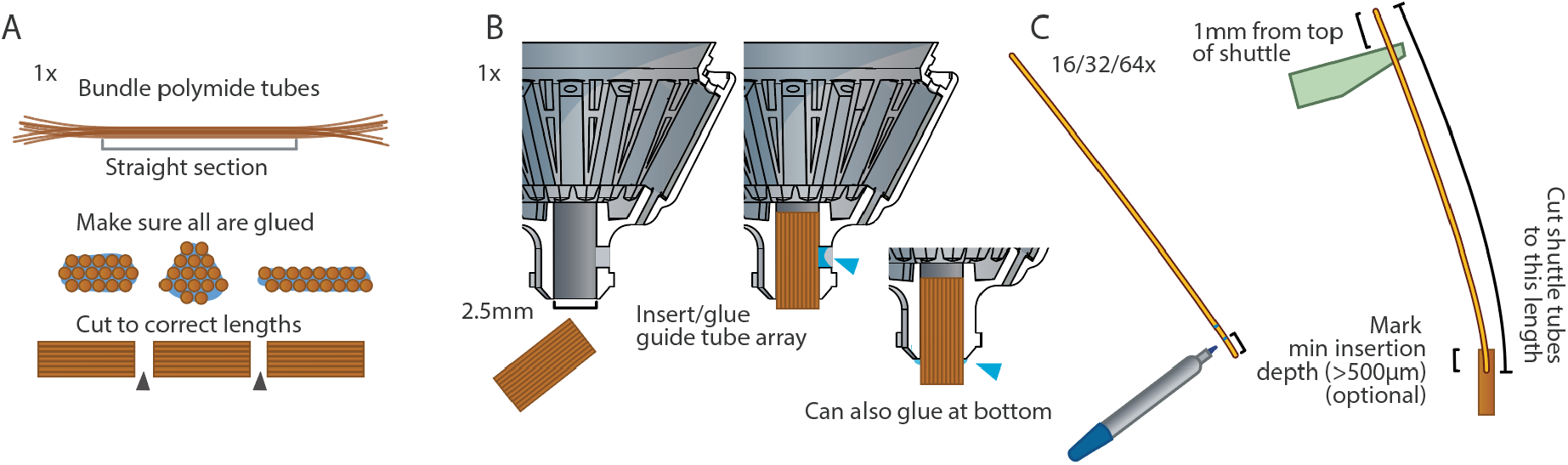
Assembly preparation. Main steps required for preparing drive assembly. **A**, Make guide tube array with the desired spacing and arrangement of polyimide guide tubes^18,34^, or use a 3D-printed or machined guide tube array. In contrast to some prior designs, the guide tube array is straight, and can therefore be prepared in one long bundle and cut to lengths later. A sufficient length of the guide tube array is crucial - see Fig. 2 and main text for details. **B**, Glue guide tube array into drive body, taking extreme care not to let glue get into the guide tubes: Insert the guide tube first, then glue with epoxy at the lateral cutout in the drive body. Alternatively, apply a small amount of glue onto the sides of the array before sliding it into the drive body, or leave a small section sticking out of the drive body and glue there. **C**, cut shuttle tubes to size. Optionally: mark minimum insertion depth so it is apparent when the tubes are properly placed later during assembly. By using a length of 1mm above the top of the shuttle (in top position) as reference, a correct length for all shuttle tubes can be determined. Now, by inserting these shuttle tubes to a depth where they extend 1mm past the shuttle (Fig. 7 step 4), a proper minimum insertion depth can be assured.

*Additional details:*

For larger bundles it can be challenging to make sure that the glue reaches the inner tubes, so building them up in layers can help. Tubes can also be coated in slow-setting epoxy before bundling to ensure proper gluing. The glued bundle can then be cut in lengths (see above) with a razorblade. If the ends of the cut pieces look squished, it can help to separate the cuts into rough and final cuts: first cut into approximate lengths, leaving half a mm or so at each end. Then cut off thin slices at the ends, holding the razorblade inwards so that the cut is straight down, and the discarded slice of polyimide tubes bends away with little effort. This procedure reduces the sideways force on the blade and can result in much nicer cuts for big arrays. After cutting, make sure that the ends of the polyimide tubes are not in any way deformed (use a fresh, very sharp blade). If any deformation is observed or suspected, the ends of the tubes can be rounded and opened slightly by carefully inserting a sewing needle to taper the opening a bit. Verify that the ends are not deformed by inserting a piece of *shuttle tube* and checking that there is not excess friction.

For making steel guide tube arrays, cutting the bundle with a wire EDM (Electric Discharge Machining) process is recommended to ensure clean edges, but almost any method that does not deform the tubes could work. The array then needs to be cleaned up with fine sand paper, and the openings need to be deburred. Small burrs left on the array could damage the polymicro tubes or tetrodes later.

For 3D-printing the guide tube array, use the provided design files as template and modify the arrangement of the guide holes while maintaining a sufficient minimal wall thickness. Different printing service providers will have different capabilities. We had good results with 200 μm holes - see section on custom parts, and the 3D files in the repository for details.

If the array is too small for the opening in the drive body, short lengths of shrink wrap tubing can be shrunk around the guide tube array, if needed in two layers, to make it large enough to press-fit into the bottom of the drive body. The array can then be secured with epoxy.

For any type of guide tube array, it is of utmost importance that the *shuttle tubes* can move through the holes with minimal friction. The amount of friction here is the main determining factor for drive mechanism hysteresis.

### Prepare drive body

The drive body, 3d-printed from our source files, should be almost completely ready for use, but in some cases modifications of the drive body are desired, such as drilling holes for guide tube arrays or other accessories. In many cases, electrical shielding should be added to the drive body. This can be done by painting the outside of the drive with conductive paint (Totalground spray paint, applied with a brush works well; a thin layer of conductive epoxy also works). Do not paint the bottom part of the drive, as this could interfere with proper glue adhesion later. Alternatively, thin tinfoil can be glued around the outside of the drive. Later, the ground wire is glued to the shield with conductive epoxy.

### Glue guide tube array to body

(This step can be swapped with the drive mechanism assembly): Insert guide tube array into the drive body, verify alignment, and glue in place (Fig. 5B). In most cases, the guide tube array should extend past the bottom of the drive body. This makes it easier to implant the guide tube array without interfering with the bone, and makes it easier to observe implant depth. This choice also depends on the desired travel distance and drive height, as gluing the guide tube array lower in the drive will equivalently increase the total implanted height of the drive. This can be especially important if the drive is to be implanted close to other implants (ground or other electrodes, head posts etc.). Inserting the guide tube array a bit lower in the drive body also reduces the bending radius of the *shuttle tubes* (Fig. 6), which improves mechanical drive function by reducing friction, and reduces the probability of breaking shuttle tubes. We therefore recommend in most cases not to extend the top of the guide tube array all the way into the drive body, and instead gluing them a bit lower.

**Figure 6:**
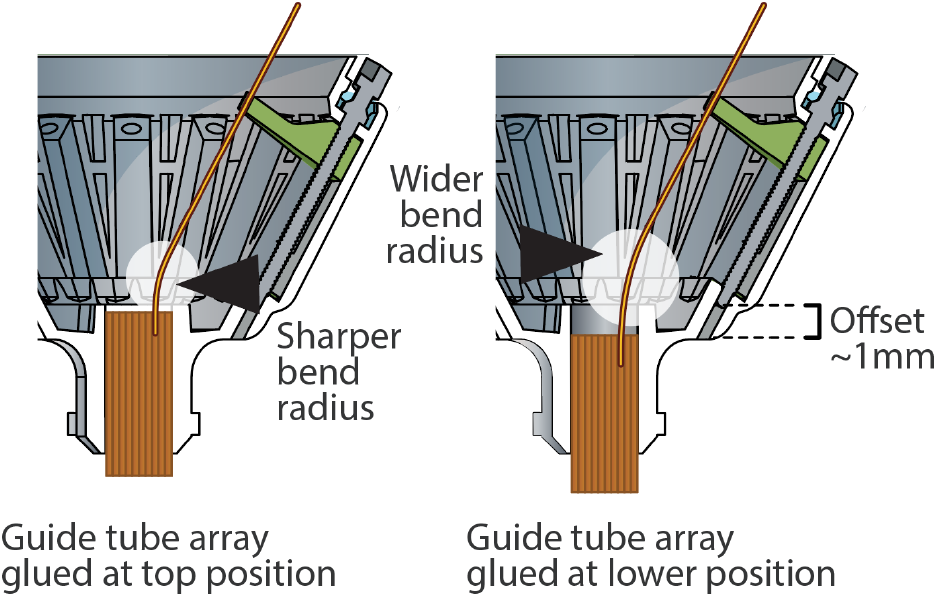
Alignment of guide tube array in drive body. Setting the *guide tube* array in the drive body with a small offset (2-5mm) reduces the bending radius of the *shuttle tubes* in lower drive positions. The effect of this offset is less pronounced on drives that do not make use of the full drive range. A separate benefit of leaving a small offset is that it makes the *guide tube* array more visible during surgery and makes it possible to fit the array into smaller craniotomies.

*Additional details:*

During insertion, the guide tube array should be held in by friction, or small pieces of tape can be used to temporarily secure it. To verify proper alignment of the guide tube array with the drive axis (or any other desired axes) it helps to insert a longer piece of polymicro tubing into one of the guide tubes to make the angle easier to measure. In the default drive body variants, an opening in the side of the guide tube array holder allows adding glue without risking glue entering the guide tubes (Fig. 5B). Once the guide tube array is stable, keep adding glue to close off any remaining openings between the drive body and the array. For this step, epoxy or thick/gel cyanoacrylate (Loctite 454 or similar) can be used.

### Assemble shuttle drive mechanisms

(This step can be swapped with the guide tube insertion step): Prepare and inspect the shuttles. If the shuttles were printed on a multi-shuttle part with some connecting material (i.e. palletized), carefully cut them apart and clean off all remains of the connecting parts. If left on the shuttles, these pieces of plastic could run into each other when the drive mechanisms are lowered. Apply a small amount of lubricant to the left and right walls of the *guide channels*; any plastics compatible grease should work here (We use Mobil Polyrex EM Polyurea Grease). The amount of grease should be small, and no grease should be allowed to get into the screw hole at the top of the *guide channels*.

*Additional details:*

For the large drive variant, two shuttle lengths are available - a shorter one to be used with guide tubes closer to the perimeter of the drive, and longer ones for guide tubes closer to the center axis of the drive. Which ones works better for which positions also depends on the angle of the guide tube arrays, and it is likely that even shorter shuttles may be advantageous for some recording geometries with angled or eccentric guide tubes.

Insert shuttles into or near the bottom positions in the *guide channel*, with the screw hole aligned vertically, and the longer protrusion at the bottom (See Fig. 7). This should require a bit of force (∼200g), so take special care not to hold the *shuttle* in a way that risks damaging the fragile hook at the tips of the shuttles. For this purpose, there is a V-shaped cutout on the top side of the shuttles (only in the smaller variant) that one side of a pair of tweezers (#3 Dumont or similar) fits into - this cutout then makes it easy to hold and press the shuttle into the guide channel without the risk of slipping.

**Figure 7:**
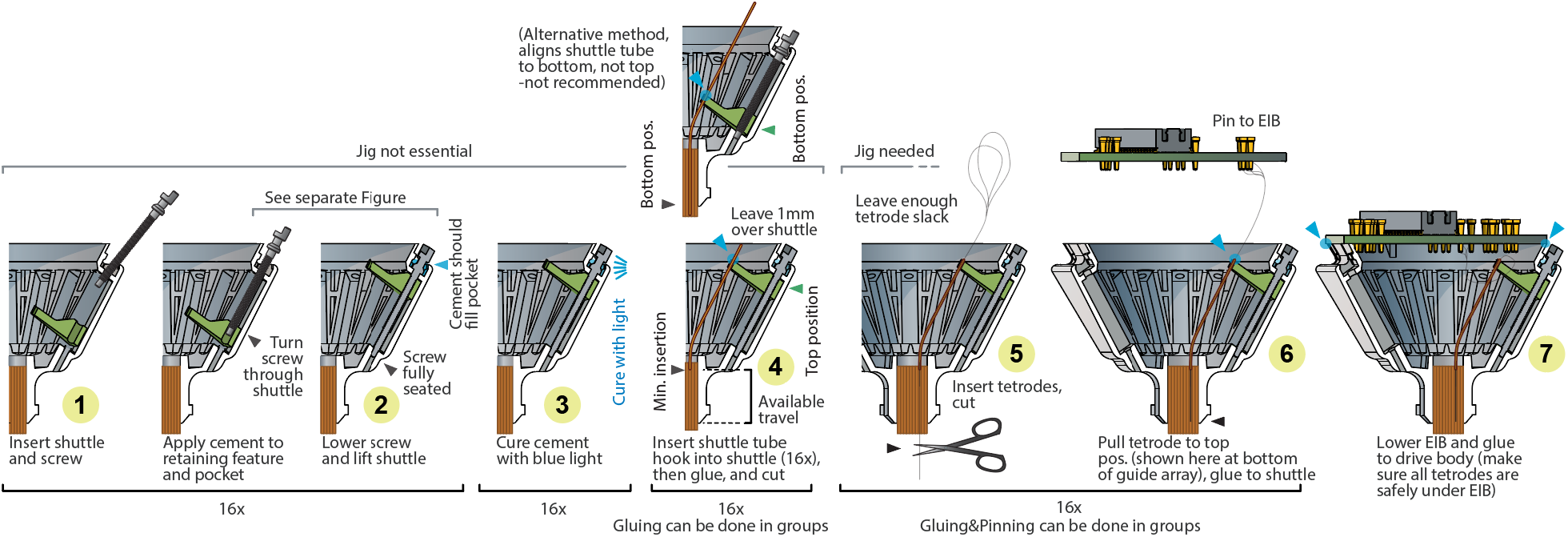
Step-by-step assembly process. This overview starts with a drive body in which the guide tube array is already inserted and glued, and ends at a loaded drive. See^18,34^ for instructions of assembling guide tube arrays. This overview omits any additional wires such as ground and/or reference, electrical stimulation, LEDs etc. Some steps can be grouped, such as inserting and seating the screws and shuttles, while others are easier if performed one drive mechanism at a time, such as loading and pinning tetrodes. See^25^ and^18^ for general procedures for tetrode assembly and loading. The procedure for the 64 drive variant is identical other than for sizing.

Insert a screw into the top of the drive body (Fig. 7 step 1) and turn it through the shuttle until the screw is seated fully in the bottom stabilizer hole and the shuttle is starting to move up (Fig. 7 step 2). Before the screw retaining collar is glued into the drive, it should be hard to lower the shuttle without exerting some down pressure on the screws. Be careful not to bend the screws, especially making sure that the shuttles are fully seated in the back of the *guide channels* so that the screws can go straight through the shuttles and into the lower guide holes. Do not apply lubricants to screws before inserting them - doing so will leave lubricant in the cavities at the top of the drive body, and interfere with gluing. Once this is done for all screws, glue the screws into the retainer pockets on the top of the drive using an appropriate glue; we recommend light cured dental cement (we use clear Vivadent Tetric EvoFlow, see git repository for details) (Fig. 7 step 3).

*Additional details:*

Curing the cement to retain the screws can be done one screw at a time, or in groups. A limiting factor to doing this in groups is that high intensity white light from the microscope illumination can cure the glue before the glue application and proper screw seating is completed. Generally, it is a good idea to work somewhat quickly if bright lights are used. See Figure 8 for details on how to properly glue the screws. Leaving some extruded cement outside of the cavities on the sides of the screw heads does not cause issues as long as the screw heads can be turned, and it is generally safer to add too much rather than too little cement.

Epoxy has been shown to not work well for retaining the drive screws, mostly because it lacks the required mechanical stiffness.

After gluing all screws into their retaining pockets, try lowering each shuttle by a little bit to ensure that the screw retention holds up - if the screw is ripped out of the pocket in this step, this indicates that either (1) the glue was not injected into the cavity properly, (2) the screw was not seated properly before curing, (3) the 3D printing quality is not sufficient to produce functioning concave retaining pockets, or (4) lubricant got into the pockets. If this happens, the mechanism can often be fixed by unscrewing the screw until the head is clear of the drive, removing the cement from the screw (gently crushing it with smooth pliers), and re-gluing the screw. Additional stability can be achieved by making sure that the glue across neighboring screws connects.

**Figure 8:**
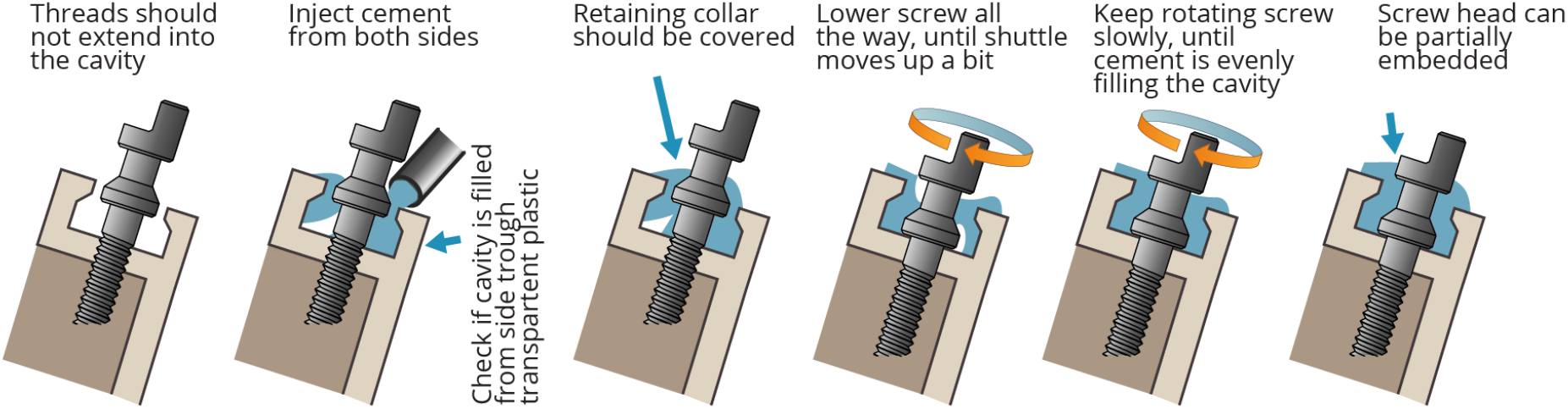
Cement application for screw retention. Step-by-step description of the process for applying light-cured cement to the screws. The main goals are to completely fill the cavity, with minimal air bubbles, and to fully cover the retaining collar of the screw. Usually some cement spills out of the cavity and extends up to the screw head; this should not cause any trouble as long as the half-moon section of the screw and a minimal section below stay accessible. The last steps of evenly covering the retaining collar of the screws can usually be accomplished by slowly turning the screw (in the direction that lifts the *shuttle* / pushes the screw into the cavity). This should pull the uncured cement around the screw, covering it. This gluing step can be done in groups of more than one screw if desired, but this can risk premature curing of the cement, especially if bright lights are used.

### Insert shuttle tubes

Move all shuttles to top position, taking care to stop as soon as there is *any* increased resistance when turning the screws, which indicates that the top of the travel range is reached. Cut pieces of polymicro tubes to the appropriate initial length (25-30mm for the rat and 15-18mm for the mouse drive, also depending on the desired travel range and guide tube array position). To determine the exact length, insert a tube into the guide tube array to its minimum insertion depth of ∼0.2 - 0.5mm and hook it into a shuttle. Now mark the tube exactly 1mm from the top of the shuttle, and cut it there. Now, by cutting all other shuttle tubes to this length and inserting them to a depth where they stick out 1mm from the shuttles, they can easily be positioned at the correct minimum insertion depth (Fig. 5C).

*Additional details:*

The minimum insertion depth is the depth of insertion of the shuttle tubes into guide tubes at which the shuttle can cleanly enter the guide tube array without major changes in curvature when pushed in. This depth depends on the *guide tube array* and relative positions of *shuttle* and *guide tubes*. An insertion of ∼0.2 - 0.5mm at *shuttle* top position has proven to work well, but more might reduce the change in *shuttle tube* curvature caused by subsequent adjustment and might yield a more linear screw to tetrode motion relationship at the expense of some travel range.

For cutting polymicro tubes, apply the same basic method as for cutting optical fibers: score with a ceramic tile, or sapphire or diamond knife, and break. The tubes can also be cut with scissors but this will result in a messy break and can cause debris to enter the tube.

Now insert *shuttle tubes* into the *guide tube array* and hook them inot the *shuttles*. They will stay there by friction alone and can be carefully moved up/down with forceps. If the tubes were cut to a precise length before(Fig. 5C), arrange them so they stick out 1mm frm the top of the shuttle to assure correct minimum insertion depth (Fig. 7 step 4).

Now glue the shuttle tubes to the shuttles with small dabs of epoxy. The epoxy should fill the opening left between the hook of the *shuttle* and the *shuttle tube* and form a small fillet around the *shuttle tube* on the top of the *shuttle*, and a layer of epoxy should be applied to the top of the *shuttle*, extending all the way into the V-shaped cutout on its top surface. Make sure that no big droplets of glue remain on the sides of the *shuttles* - these can touch neighboring *shuttles* and cause issues at the bottom of the travel range. Extra epoxy on the top surface should cause no problems.

Now cut the *shuttle tubes* as close to the *shuttles* as possible, ∼0.1 mm above the *shuttles*. Leaving more length here increases the risk that the shuttle tubes will collide with a gold pin later when the EIB is lowered. This cut can be made with scissors, which will leave a more messy break, that has not proven to cause significant issues in practice, or with a diamond/sapphire/ceramic knife with the score and bend/pull method, which will leave a cleaner cut but may be a little harder to do quickly.

*Additional details:*

Alternative method (recommended for the rat version, but also useful for the mouse version)- leave *shuttles* at top position. Insert and push down the *shuttle tubes* to the bottom end of the travel range. This can be verified by observing the *shuttle tubes* are visible at the bottom opening of the guide tube array, but not protruding, or by pushing them down against a surface that is flush with the guide tube array bottom. The upside of this method is that the bottom position is easily verified and it is easy to achieve the maximum travel range. Now place the *shuttle tubes* into the hooks in the *shuttles* and pull them up by the desired travel range, by using a small ruler or calipers. Now cut them to a common length, and glue them to the hooks.

In either case it is not recommended to place *shuttle tubes* into the hooks of the *shuttles* while in their bottom position, as this places considerable strain on the hooks and can break them. Once the *shuttle tubes* are glued to the *shuttles*, the bending moment is supported by the epoxy, not the hooks, so lowering them after gluing does not have this risk.

The most important parameter in this step is that the *shuttle tubes* must be able to move into the *guide tubes* with minimal friction. The main cause of friction is that the guide tubes can be left in a slightly oval shape after cutting the *guide tube array*, in some case barely enough to cause problems but not be immediately obvious visually. In this case, try opening up the ends of the guide tubes by gently inserting a sewing needle (not a beveled needle) until the opening is very slightly flared and rounded.

Load tetrodes: Make the desired number of tetrodes^22,26,35^. For maximum build speed, use a high-speed twister^24^. Insert the drive body into the jig (if not done before) and position the EIB ∼10mm (mouse) and ∼20-30mm (rat) from the drive body. For the mouse drive, the distance needs to be >15mm to allow rotation of the EIB later, and <20mm space to avoid tetrode loops sticking out of the drive later. This distance is chosen to be long enough to allow inserting and gold-pinning tetrodes into the EIB, and leave enough slack in the tetrodes for later drive motion (the EIB will later be rotated by 180° or 360° later in order to bring tetrodes under the EIB and avoid them from sticking out). Now the tetrodes can be inserted into the *shuttle tubes*, cut, glued in place (Fig. 7 step 5), and connected to the EIB (Fig. 7 step 6). Inserting the tetrodes can be a bit harder than in traditional drives with polyimide (not polymicro) tubes if the cuts in the polymicro *shuttle tubes* are not clean, like if they were cut with scissors, but it should be relatively easy with some practice.

*Additional details:*

There are different possible orders of steps in loading the drives. The position of the shuttles (top vs. bottom) and order (pinning tetrodes individually after loading or after loading all) can be chosen according to specific requirements and preferences.

With *shuttle* in top position, insert tetrode into *shuttle tube*, and pull through until branching point is slightly below EIB. Make sure that tetrode is at correct length to allow enough slack under the EIB once lowered. This depends on whether you want to rotate the EIB during lowering, and by how much.

Now perform a final cut of the tetrode^18,25^; verify cut with microscope if desired^12^, and pull the tetrode back until it is flush or slightly recessed relative to bottom of *guide tube array* (or leave it sticking out, depending on application), verify this with microscope. Glue tetrode to *shuttle tube* with epoxy.

*Additional details:*

Alternatively tetrodes can be left in this position, and gluing can be deferred until after gold-plating, if the option of replacing tetrodes later is desired. Now cut tetrode loop, or trim non-fused end of tetrode (ideally in a way that leaves the 4 ends at different lengths) and insert into EIB holes and pin. With experience, the steps leading up to gluing the tetrodes can be done in groups of 4 or more tetrodes in order to reduce the numbers of batches of epoxy and speed up the process. Depending on the length of the tetrodes, and how high the EIB is held, the tetrodes will be bending significantly and sticking out under the EIB at this stage.

Load all tetrodes, then pin tetrodes: Tetrodes can also be loaded first, and then pinned to the EIB in one go. For this method, tetrodes can be loaded without the EIB in the jig. Tetrodes are then cut (and glued to the shuttles) left sticking out the shuttle tubes, with 15-20mm of length between the top of the drive and the branching point of the tetrodes (mouse drive) or 30-40mm (rat drive). This length must be present in order to allow rotation of the EIB by 360° (or at least 180°) during lowering of the EIB (Fig. 9). It is best to verify the length of the tetrodes with the EIB and loading procedure before proceeding with all tetrodes. Alternatively, gluing of the tetrodes to shuttles can also be performed after pinning to the EIB, but this requires extra caution to make sure that the zero position of the tetrodes is precisely known.

**Figure 9:**
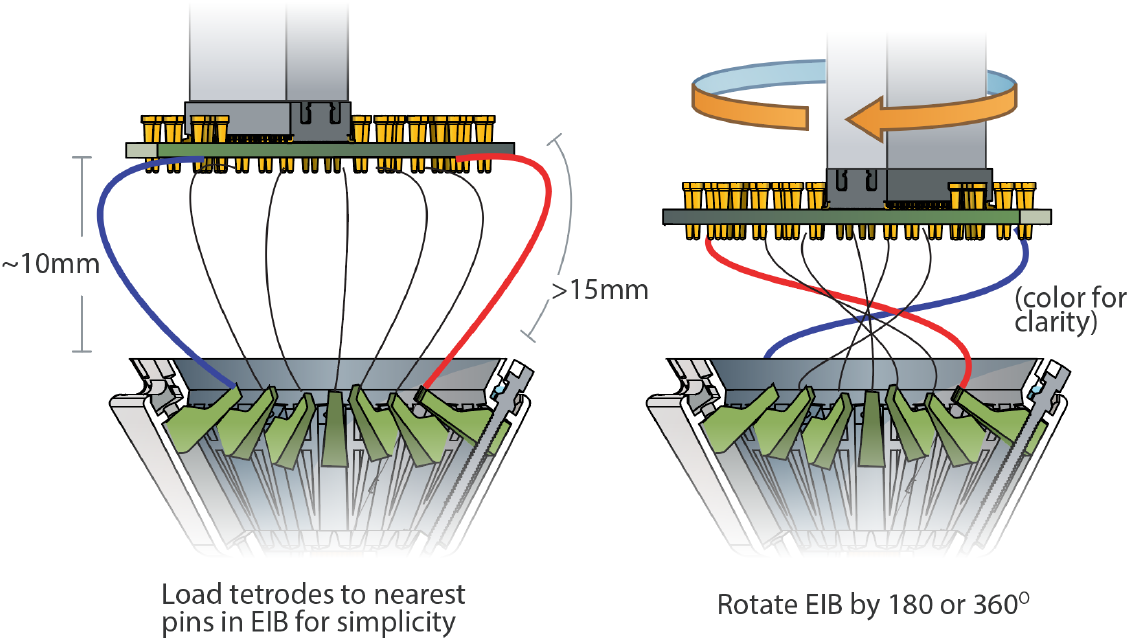
Rotating EIB before lowering to bundle up tetrode wires. Rotating the EIB by 180° or 360° while lowering it tucks the tetrodes in under the EIB, making it easier to avoid accidentally gluing tetrodes to the rim of the EIB. Mouse: distance 10-12mm, tetrodes 15-20mm, Rat: distance 20-30mm, tetrodes 30-40mm. Test these distances with a few tetrodes first before loading entire drive.

It is also possible to load two tetrodes into individual *shuttle tubes*. This method can be useful for targeting small regions.

If desired, the tetrodes can be gold-plated now, before gluing them to the *shuttle tubes* or lowering the EIB, using a plating apperatus that fits into the assembly jig (see git repository at https://github.com/open-ephys/shuttle-drive). This ordering of steps makes it easy to replace tetrodes that are found to be faulty during the plating step.

When lifting or lowering the *shuttles* to the top and bottom ends of the travel range, extreme care needs to be taken to avoid crashing the *shuttle* into the ends of the *guide channel*, which can strip the threads in the *shuttles*. It is possible to feel the *shuttle* reaching the end of the travel range by a small increase in the force required to turn the screw, but this method is fairly risky because continued rotation of the screw even by a bit after this point will strip the threads.

Once all tetrodes are glued to the *shuttles*, lower the EIB onto the drive body by adjusting the linear slide on the assembly jig, and glue in place with epoxy (Fig. 7 step 8). While lowering, take extreme care not to leave tetrode wires sticking outside from under the EIB. In some cases, if a lot of loose tetrode wire is left under the EIB, leftover wire needs to be manually pushed under the EIB. Ideally, this is done with a smooth tool such as a blunt needle or smooth (carbon/ceramic tipped) forceps to avoid accidental snagging of tetodes. To ensure that all tetrodes are safely under the EIB, the EIB can be rotated by 180° or 360° before or during lowering (Fig. 9). This requires leaving sufficient slack in the tetrodes, around 15-20mm for the mouse and 30-40mm for the rat version work well, but verify using one or two tetrodes before loading entire drives for this method. Epoxy should be applied on the outside edge of the EIB once it is lowered, not under the EIB, to avoid gluing tetrodes to the EIB by accident.

After gold plating, the gold pins should be secured with a small amount of epoxy. A thin layer of epoxy should extend from the sides of the drive body, over the edge of the EIB and up to the gold pins to ensure a stable mechanical connection.

*Additional details:*

If using the upcoming Open Ephys headstages (Newman, Zhang et al., in prep), make sure throughout this step that no piece of wire, epoxy or anything else extends above the level of the gold pins, approximately 0.6mm from the surface of the EIB PCB. Any protrusions here will make it impossible to connect the headstage later. If there are pieces of glue or even wire or gold pins extending past this limit, they can be filed down with fine sandpaper.

### Shield drive and add ground wire connection

For shielding and grounding, two connections need to be made: (1) Ground (GND) on the EIB needs to be connected to a ground screw or wire on the animal, often with a removable connector, such as a single mill-max pin or similar. In most cases the amplifiers reference input (REF), which provides a reference with higher input impedance (though on Intan chips not the same as the other inputs, see Intan data sheets for details), can be tied to GND. (2) GND need to be connected to a shield/faraday cage that surrounds the drive. For the latter, conductive paint can be applied to the drive body and connected to the ground wire (Fig. 10), or directly to the EIB with conductive paint.

**Figure 10:**
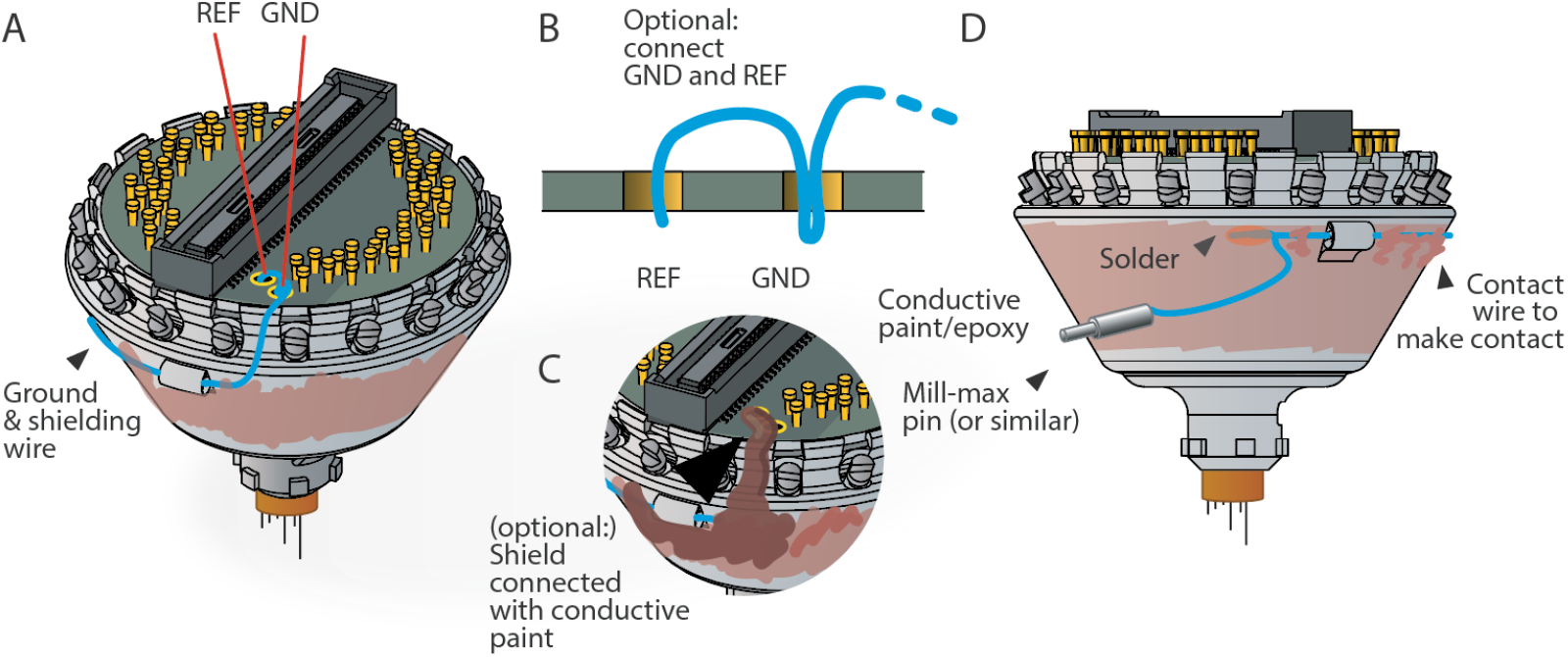
Shielding the drive and connecting the ground wire. In almost all applications the drive should be shielded. **A**, schematic of the recommended shielding scheme. Here, GND and REF are shown connected. This is not always the ideal configuration, and some use cases require separate GND and REF connections, but works well for most applications. The ground wire (blue) is soldered to GDN, or GND and REF, and threaded through the loop at the side of the drive, making sure to place it between the screws in a position that allows screw function. The wire is then laid along the side of the drive. **B**, recommended schema for connecting GND and REF with one wire. **C**, The shield can also be connected to GND by painting a trace up to a GND via on the EIB with conductive pain or epoxy. **D**, the ground wire can be laid flat along the side of the drive, threaded through the second loop, and then used to solder to the wire for the ground screw connection. The drive can then be painted with conductive paint to form a shield, or aluminum foil can be lgued to the drive and connected to GND with conductive paint.

An easy and reliable way to construct the shield is to use a high quality 2-part conductive epoxy (See repository at https://github.com/open-ephys/shuttle-drive for part numbers) to paint the shield and connect it to the ground wire. This results in a robust shield that will easily bond with further glue during surgery (for attaching the ground connector etc.). We recommend that the shield is not extended all the way to the bottom of the drive (Fig. 10), so that during implantation, the cement used to attach the drive can bond to the bare plastic of the drive body.

*Additional details:*

Alternatively, conductive paint can just be applied to the side of the drive, and a strip of paint extended up to a GND and/or REF via on the EIB, ommitting the ground wire connection to the shield. A ground wire, soldered to the appropriate vias would then connect to the animal ground without connecting to the shield.

Now the drive is ready for gold plating and implantation - see Nguyen et al. 2009^25^ for overview of the method, and Ferguson et al. 2009^12^ and Keefer et al. 2008^17^ for methods for even lower (< 100kΩ) impedance plating.

## Part availability and manufacturing

The parts of the presented drive design can be ordered from a number of sources and assembled easily with common tools, though a few specialized tools are needed. Most of the wearing parts are 3D printed and can be acquired with low minimum order quantities and lead times. For a list of suppliers and part numbers please refer to the documentation on the repository (https://github.com/open-ephys/shuttle-drive), and please submit additions as pull requests if availability and suppliers change.

### Assembly jig

Most 3D printed parts here can be made with medium resolution 3D printing, possibly even with well tuned fused plastic deposition printers. Other components are all common stock parts (Screws and precision dowels). See the git repository for the recommended print processes for all parts.

### Screwdriver for drive screws

A custom screwdriver (Fig. 11) is needed to turn the screws during construction and for later adjustment. Such screwdrivers can be made by gluing or soldering a screw into the end of a piece of steel tube with an ID of close to 1mm. Thin-walled tubes are recommended because thicker tubes will interfere with the rest of the drive body when adjusting. The screw used for this should be inspected to make sure that the protruding half-moon section is on the smaller side, so it can interface with other screws. It can also help to carefully bevel the edge of this section with fine sandpaper. The screw should be slightly recessed into the steel tube so that the screwdriver can be locked onto a screw axially first, and then rotated to snap into the mating screw rotationally. A 3d-printed screwdriver handle and detailed tube measurements and sources are available on the git repository.

**Figure 11:**
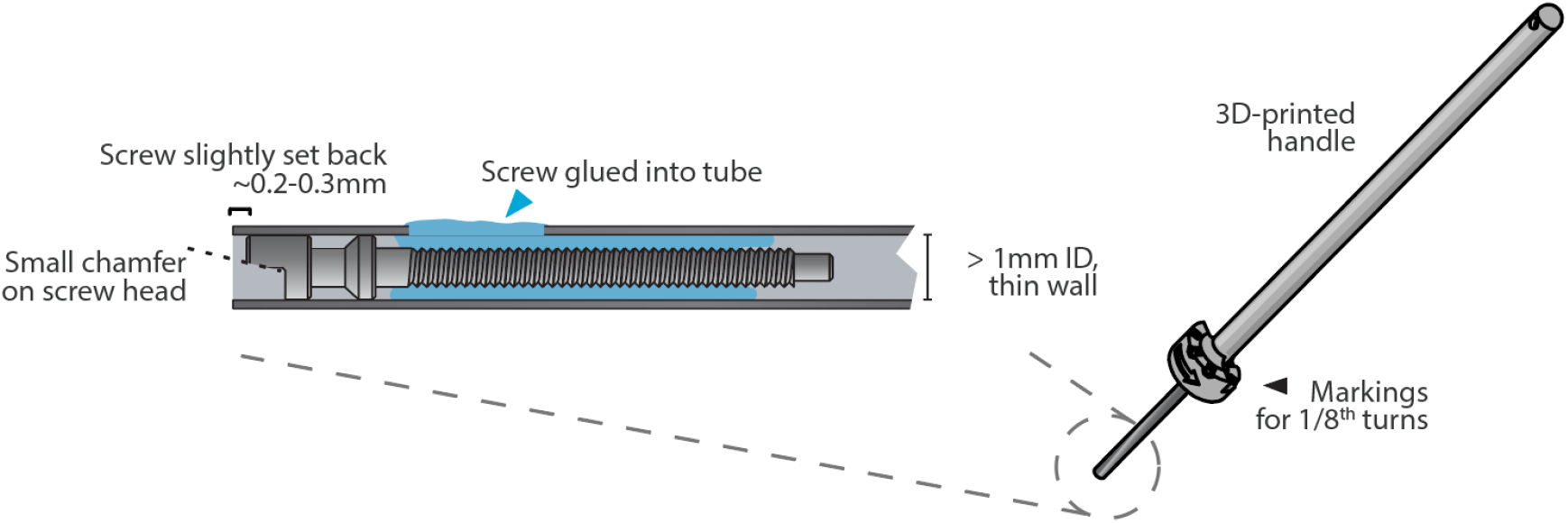
Custom screw driver for tetrode adjustment. Close up of the custom screw driver, made from a stainless steel tube with an ID that snugly fits the screw head. A handle can be made by applying heat-shrink tubing to the cannula, or by 3-D printing the handle from files available on the git repository.

### Drive shuttles

Require very high resolution 3D printing in order for the screw hole tolerances to be stable, and to allow printing of the guide hole for the shuttle tube (∼300μm hole). We use Microfine Green (Fine Line Prototyping / Protolabs, 85 Shore D hardness), with 0.0254mm layer thickness and 0.0508mm x/y feature size. Different printers or printing processes will very likely require changes in the drive shuttle width (the two sides that press against the guide channel). For this purpose, use a test part design (see the repository) that combines various different sizes in small increments in one part, and choose the width that fits best: The guide channels on the drive bodies will not all be the same width due to printing process variability. The shuttles should require some force to push into the tightest of the guide channels, but should still move up easily when the screw is turned (prior to gluing the screw retaining collar) without the screw causing damage to the top of the drive body. In the loosest channels, the shuttles should move up and down without or only with minimal visible sideways motion. Before the screw is glued, it should be hard to lower the shuttle without exerting some down pressure on the screws. Many 3D printing suppliers will require many shuttles to be stuck together in one part for easier printing and cleanup, this is called palletizing. Versions with palletized shuttles are included in the repository but might have to be edited for different suppliers.

### Shuttle tubes

We use Molex polymicro tubing (polyimide covered fused silica tubing), ID 100±4μm, OD 164±06μm, CT 12μm-part number TSP100170. Replacing this part with pure polyimide tubes will likely cause problems during drive adjustments because pure polyimide is not stiff enough to withstand the bending forces that occur when lowering the drives.

### Printed guide tube array

(optional - we recommend using polyimide tubes): these require very high resolution 3D printing, we used Microfine Green for initial tests, as a 4.8mm straight extrusion with holes printed as hexagons with a 0.19mm inscribed diameter, in a hexagonal pattern with 0.33mm pitch, resulting in uniform 0.14mm walls between the holes. The guide tube array could also be CNC machined in delrin/polyimide type plastics, but these would require significantly high aspect ratio drilling. We have not tested this approach extensively enough to comment on stability, biocompatibility, and use with solvents and lubricants.

### Polyimide guide tube array

These can be made the traditional way^18^ from bundled Polyimide tubes. For the ∼160μm OD shuttles tubes we use 30ga tubes with ID: 180μm OD: 295μm. Larger, tubes could also be used to increase the tetrode pitch, or thinner walled tubes to decrease it. We recommend a minimum OD of 250μm as a safe closest pitch - closer spacing requires extra caution to avoid tissue damage.

### Drive body

This part requires high resolution 3D printing in order for the sides of the guide channels to be smooth enough for good drive function. We use 3D Systems Accura 60 (SLA) material. We also previously used Somos WaterShed XC 11122 with acceptable results before. The drive bodies are printed with with 0.0508mm layer thickness and 0.1016mm x/y feature size. For best consistence between the guide channels, orienting the drives so that the printed layers are orthogonal to the vertical axis of the drive is recommended. Otherwise, the different guide channels are printed at different angles and may end up with different widths and surface textures which makes fine-tuning their width difficult. Finally, for printing processes that require a support structure, we recommend orienting the drive so that this support structure attaches to the outside of the drive.

### Drive screws

Require high quality custom screw production, we recommend using a trusted vendor, as not all makers of micro screws can provide the required high surface finish quality especially on a M0.6 screw. Typical minimum orders are 500-1000 screws, and lead times are 2 months. The repository at https://github.com/open-ephys/shuttle-drive and Open Ephys online forums should have information on vendors.

### EIBs

Require medium quality PCB manufacturing (4 layers, 5mil trace/space), and some soldering, these can be assembled by hand. The main constraint is that the hole size for the vias needs to accommodate small Neuralynx gold pins, so 300μm ±15% vias are needed. The 64 channel omnetics EIB requires good alignment between the two 32 channel omnetics connectors and hand-soldering these requires some experience. The correct spacing between the connectors (1mm gap for the Intan 64 channel headstage) can be maintained by clamping the connectors to a spacer during manual or reflow-soldering. Extra attention is needed for the OD of the EIBs. The outer dimensions have to remain under 15.95mm OD. If too large, the EIB can get in the way of accessing the screws for adjusting. If this is an issue, the EIBs can easily be sanded down a bit.

## Part sizing tolerances

Some parts of the drive design require close tolerances. Here is a partial list of the most important places where parts need to fit together in a specific way. If changes in any of the affected parts are made, or even when changing suppliers and/or manufacturing methods, these tolerances need to be verified. This is especially true for changes in 3D-printing or PCB suppliers.

### EIB OD

The OD of the EIB (15.94mm for the mouse version) must fit into the retaining ring feature on the drive body. If this is not the case, the EIB can be sanded down. In some cases it might be acceptable to glue the EIB on top of the retaining ring. For the rat version, additional cutouts on the EIB are required to fit the drive body.

### Guide tube ID

Needs to fit polymicro shuttle tube OD of ∼170 μm. See section ‘Insert shuttle tubes’ for discussion of shuttle tube fit.

### Shuttle width vs. guide channel width

The shuttles need to fit the guide channels. Nominal channel and shuttle widths are channel: 1.14mm; shuttle: 1.15mm for mice, and channel: 1.2mm; shuttle: 1.14mm for rats. Printed sizes differ somewhat based on printing process, and adjustments might be required. The drive body channels are expected to have more variability than the shuttles due to the lower spec printing process. A fit that is too tight manifests in shuttles that are hard to insert into the channels by hand, and in screws breaking through the top of the drive body when attempting to raise a drive mechanism. A fit that is too loose manifests in sideways motion of the shuttle when screws are turned. A little bit of sideways motion during adjustment should not cause any issues. During building, it is important to feel for the amount of force required to insert the *shuttles* into the *guide channels*. If the force is too large (> 400g), one or two passes of the shuttle over fine sandpaper can be used to make the shuttle fit. If this occurs systematically, adjusting the channel or shuttle width (whatever is more economically viable) is recommended.

### Guide channel smoothness

Guide channel walls need to be smooth enough to allow for easy shuttle motion. If prominent steps/ridges from the 3D-printing can be felt with forceps, a higher quality printing method, or a different post-print polishing protocol, should be used.

### Screw hole ID (top and bottom)

The hole IDs for the drive screws at the top and bottom of the drive mechanisms are not particularly critical. Because the section of the screw that sits in the upper hole is smooth, as long as screws can fit through the top they are not too small, even if the threads of the screws catch on the holes. Conversely, a bit of play is acceptable, as long as the screws are not pulled down through the holes when attempting to raise a drive shuttle.

### EIB envelope and height with gold pins

The next-gen headstage sits flat on top pf the EIB (Newman, Zhang et al., in prep). When modifying the EIB, or soldering additional ground wires etc. make sure that the height of the gold pins and any other added material does not interfere with the headstage. The overall free height over the PCB surface on the mouse version is ∼1.0 mm, though some areas allow a higher tolerance.

## Discussion

Our drive design provides a combination of high performance in terms of low weight, high drive count and robustness, and ease of assembly. The major limitations imposed by these design priorities are the use of completely custom screws, and the need for high quality 3D printing. However, most existing designs of comparable performance (weight and drive count), particularly the flexDrive^34^, share these downsides, while requiring much longer, more complicated assembly.

This design significantly reduces the effort required to build drives such that drives can be completed in less than a day. This makes it viable to employ drives in ways that were previously too costly. For example, it is now viable to implant entire cohorts of animals before a task training period even if not all of the mice end up learning the task. Currently, in such circumstances only trained mice might have been implanted to reduce the number of required drives, at the cost of having to interrupt the behavioral task training with a surgery and recovery period. This improvement is compounded by the speed of modern tetrode-twisting tools^24^. The tetrode loading step, which requires individual insertion, cutting, and gold-pinning of tetrodes now remains as the most time-consuming step by a large margin, and improvements in this procedure will further bring down the effort required to build tetrode drive implants.

The presented design can be easily modified to accommodate different recording targets, numbers of drives, and other attachments. Most of these changes only require modification of one 3D printed part and are relatively simple. The basic building block, a guide channel with a screw guide and retaining pocket, together with the custom screws and printed shuttles can be translated to other applications that require precise and small linear adjustment mechanisms.

All design files, documentation, and this manuscript are available on Github at https://github.com/open-ephys/shuttle-drive and are licensed under the CERN OHL v. 1.2. See the git repository for a copy of the license. This means that if modifications are made and distributed, the modified version of the project must carry the same license conditions, which ensures that the whole community will continue benefiting from improvements^31^. We particularly encourage improvements to be shared as part of the existing repository, or as forks, in order to make them easily discoverable.

## Acknowledgments

We thank Marie-Sophie Helene van der Goes, Lukas Fischer, Dimitra Vardalaki, Hector Penagos, Josefina Correa Menendez, Ksenia Nikolaeva, Nolan Garskovas, Stefan Oline, Chris Angeloni, Priyanka Gupta, Liora Las, Ayelet Sarel and Nachum Ulanovsky for helpful discussions and comments on the drive building process, Filipe Carvalho and Lidia Fortunato for help with the design, documentation and manufacturing, and Quique Toloza for helpful comments on the manuscript.

